# CD73 generated extracellular adenosine promotes resolution of neutrophil-mediated tissue injury and restrains metaplasia in pancreatitis

**DOI:** 10.1101/2022.09.17.508367

**Authors:** Baylee J. O’Brien, Erika Y. Faraoni, Lincoln N. Strickland, Zhibo Ma, Victoria Mota, Samantha Mota, Xuebo Chen, Tingting Mills, Holger K. Eltzschig, Kathleen E. DelGiorno, Jennifer M. Bailey-Lundberg

**Affiliations:** Center for Perioperative Medicine, Department of Anesthesiology, McGovern Medical School, The University of Texas Health Science Center at Houston, Houston, TX, 77030; Gene Expression Laboratory, The Salk Institute for Biological Sciences, San Diego, CA, 92037; The Graduate School of Biomedical Sciences, The University of Texas MD Anderson Cancer Center and The University of Texas Health Science Center at Houston, Houston, TX, 77030; Department of Biochemistry, McGovern Medical School, The University of Texas Health Science Center at Houston, Houston, TX, 77030; Department of Cell and Developmental Biology, Vanderbilt University, Nashville, TN, 37232

**Keywords:** CD73, acinar-to-ductal metaplasia, inflammation, purinergic signaling

## Abstract

**Background and Aims:** Pancreatitis is currently the leading cause of gastrointestinal hospitalizations in the US. This condition occurs in response to abdominal injury, gallstones, chronic alcohol consumption or, less frequently, the cause remains idiopathic. CD73 is a cell surface ecto-5’-nucleotidase that generates extracellular adenosine, which can contribute to resolution of inflammation by binding adenosine receptors on infiltrating immune cells. *We hypothesized genetic deletion of CD73 would result in more severe pancreatitis due to decreased generation of extracellular adenosine*.

**Methods:** CD73 knockout (*CD73*^*-/-*^) and C57BL/6 (wild type, WT) mice were used to evaluate the progression and response of caerulein-induced acute and chronic pancreatitis.

**Results:** In response to caerulein-mediated chronic or acute pancreatitis, WT mice display resolution of pancreatitis at earlier timepoints than *CD73*^*-/-*^ mice. Using immunohistochemistry and analysis of single cell RNA-seq (scRNA-seq) data, we determined CD73 localization in chronic pancreatitis is primarily observed in mucin/ductal cell populations and immune cells. In murine pancreata challenged with caerulein to induce acute pancreatitis, we compared *CD73*^*-/-*^ to WT mice and observed a significant infiltration of Ly6G+, MPO+, and Granzyme B+ cells in *CD73*^*-/-*^ compared to WT pancreata and we quantified a significant increase in acinar-to-ductal metaplasia demonstrating sustained metaplasia and inflammation in *CD73*^*-/-*^ mice. Using neutrophil depletion in *CD73*^*-/-*^ mice, we show neutrophil depletion significantly reduces metaplasia defined by CK19+ cells per field and significantly reduces acute pancreatitis.

**Conclusions:** These data identify CD73 agonists as a potential therapeutic strategy for patients with acute and chronic pancreatitis as adenosine generation and activation of adenosine receptors is critical to resolve persistent inflammation in the pancreas.

**Graphical Abstract:** 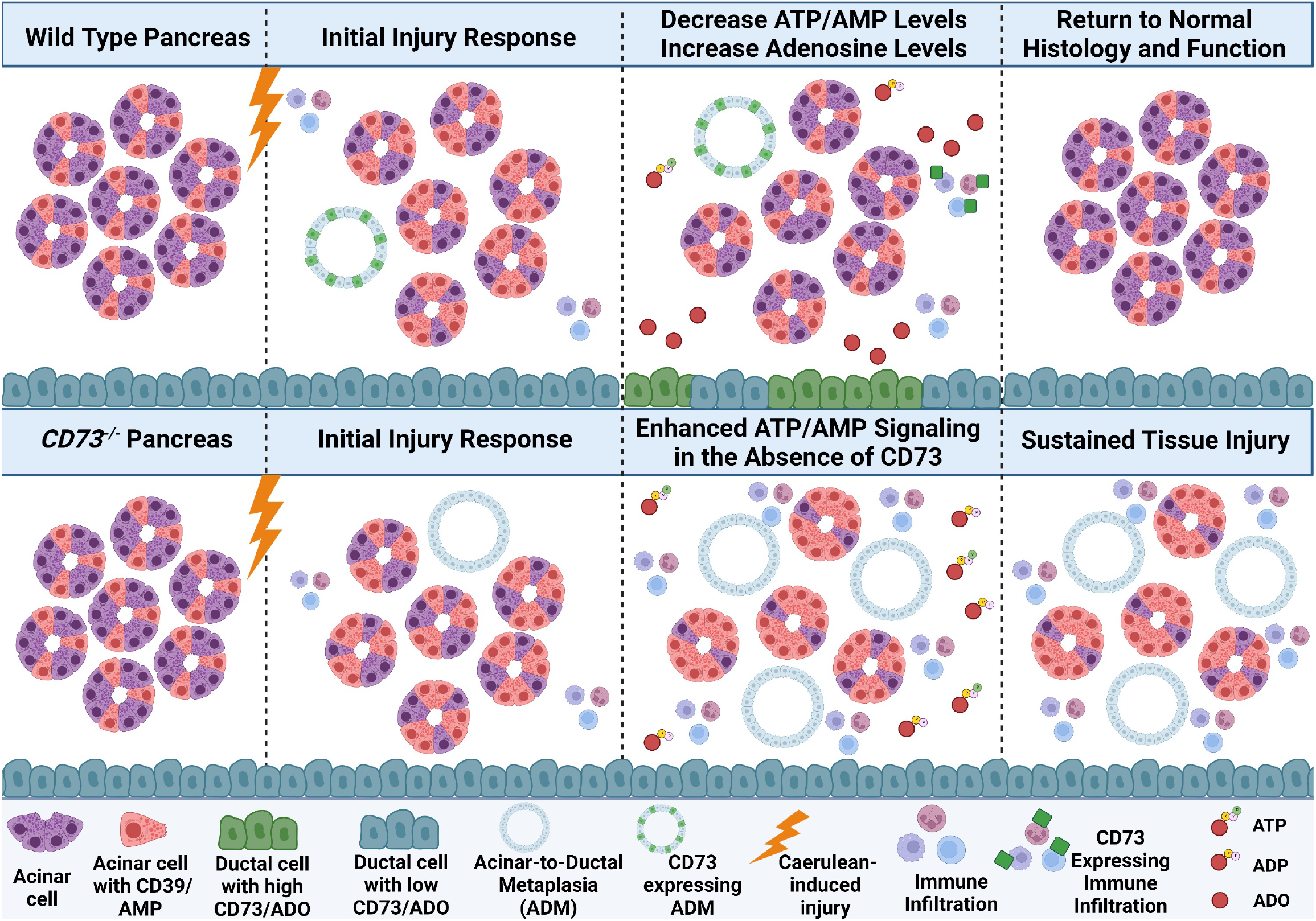

## INTRODUCTION

Pancreatitis is an inflammatory condition of the pancreas characterized by severe abdominal pain [1]. The incidence and morbidity of pancreatitis has increased significantly over the past few decades and acute pancreatitis is now the leading cause of gastrointestinal diagnosis for inpatient hospitalizations in the United States [1, 2]. This condition occurs in response to chronic alcohol consumption, gallstones, abdominal injury, or, less frequently, the cause remains idiopathic [3]. In alcohol-associated pancreatitis, alcohol induces the aberrant intracellular activation of trypsin within the pancreatic acini resulting in autodigestion of the organ [4]. Additionally, alcohol releases the secretagogue cholecystokinin (CCK) from duodenal cells that stimulates secretion of zymogen granules in pancreatic acinar cells resulting in a profound systemic inflammatory response, acinar-to-ductal metaplasia (ADM), and fibrosis [4]. Animal models of acute and chronic pancreatitis have been developed to study mechanisms of acinar cell responses to injury by administering supraphysiological concentrations of caerulein, a CCK analog [2].

Adenosine (ADO) has been recognized for decades as an important physiologic and pharmacologic regulator that signals through cell surface receptors to regulate cell function [5, 6]. More recently, adenosine has been described as paracrine regulator of the tumor microenvironment (TME) and immunosuppressive metabolite elevated in a number of solid tumors including pancreatic ductal adenocarcinoma (PDAC) [7-15]. While inhibitors targeting CD73 or adenosine receptors are therapeutic targets for PDAC patients [16], under acute or chronic inflammatory conditions, adenosine can promote fibrosis or reduce inflammation, both critical components of wound healing and repair necessary for tissue regeneration [17-25]. Adenosine triphosphate (ATP) is the primary source of energy for cellular processes localized in the intracellular space; however, in response to inflammation or hypoxia, ATP is released into the extracellular space and has been shown to promote pancreatitis [26, 27]. In the presence of ectonucleotidase triphosphate diphosphohydrolase-1, NTPDase1, (CD39), extracellular ATP is rapidly converted to adenosine diphosphate (ADP) and monophosphate (AMP), which is subsequently converted to adenosine by a cell surface ecto-5’-nucleotidase, CD73 [28-30]. Generation of extracellular adenosine by CD73 and stimulation of adenosine receptors has been shown to promote fibrosis by signaling through A_2A_ and A_2B_ receptors in the lung, skin and liver, however, the A_2B_ receptor restrains fibrosis in the heart [31]. In contrast, adenosine can also promote resolution of inflammation by inhibiting the inflammatory function of neutrophils by signaling at higher concentrations via A_1_ or A_2_ receptors to prevent tissue injury from prolonged inflammatory responses [25, 29, 32, 33]. The adenosine ENT1 transporter facilitates the uptake of extracellular adenosine across the cell membrane and is one mechanism to downregulate extracellular concentrations of adenosine [34]. Adenosine signals via G-protein coupled receptors: A_1_, A_2A_, A_2B_, and A_3_ [35-38]. The A_1_ G_i_-coupled receptor has a high affinity for adenosine while the A_2A_ and A_2B_ G_s_-coupled receptors have a lower affinity [35, 39-41]. Therefore, at early stages of inflammation, low local concentrations of adenosine promote immune recruitment via the A_1_ receptor while later high adenosine concentrations can suppress immune activity via the A_2A_ and A_2B_ receptors [35, 42].

Adenosine binding to adenosine receptors modulates both the innate and the adaptive immune response to hypoxia, inflammation, and tissue repair [30]. The A_2A_ and A_2B_ receptors exert anti-inflammatory effects by inhibiting neutrophil chemotaxis, attachment to vascular endothelial cells, and phagocytosis [14, 35, 43, 44]. Inflammatory macrophages are inhibited through A_2A_ and A_2B_ activation resulting in decreased production of cytokines including IL-1β, IL-18, IL-6, and TNF-α [14, 29]. Additionally, CD4+ T cell activation and proliferation and natural killer cell cytotoxic functions are inhibited by A_2A_ receptor activation [14]. In contrast, decreased activity of CD73 and extracellular adenosine are associated with amplified activation and chemotactic functions of immune cell populations [29]. The purinergic P2X and P2Y family of receptors are expressed on neutrophils and promote neutrophil-mediated oxidative burst-induced tissue injury in the presence of ATP [45].

In response to injury, pancreatic acini can undergo acinar-to-ductal metaplasia (ADM), a metaplastic event that limits pancreatic tissue damage via a rapid decline in zymogen production [46-48]. Experimental studies have shown injured acinar cells activate a shift in gene expression regulated by Mist1 and Ptf1α to transdifferentiate away from their specified cell type and function, which consists of highly specialized cells involved in the production and secretion of digestive enzymes, towards a ductal phenotype [44, 49]. ADM trans-differentiation is also triggered by innate and pro-inflammatory immune cells, including neutrophils and macrophages, that infiltrate the pancreas resulting in elevated secretion of inflammatory cytokines including RANTES and tumor necrosis factor α (TNF)[50]. Differentiation into a cell type with ductal characteristics demonstrates a pancreatic repair process under strong positive selection in pancreatitis [23, 26]. These mucinous populations can subsequently seed tuft cell and enteroendocrine lineages as further reparative mechanisms [51].

## RESULTS

### CD73 is expressed on ductal cells in human chronic pancreatitis

Recent literature has described that CD73 is elevated in PDAC epithelial cells resulting in adenosine generation and an immune-suppressive tumor microenvironment [9-11]. In the normal pancreas, CD73 is expressed in vascular cells, but limited to no expression is observed in acinar or ductal cells. In contrast, NTPdase1 (CD39) is expressed in acinar cells and blood vessels, which have ATPase activity; yet CD39 is expressed at low levels in normal pancreatic ducts [52, 53]. As chronic pancreatitis is a risk factor for development of PDAC [54], and divergent roles for adenosine have been described in inflammatory diseases, we wanted to investigate the cell-type specific localization of CD73 in human and murine chronic pancreatitis and determine if CD73 is an important determinant of pancreatitis severity. To evaluate cellularity of CD73 expression, we analyzed previously published single-nucleus RNA-sequencing data (sNuc-seq) generated from two patients with chronic pancreatitis (totaling 2726 nuclei)[51]. These data revealed NT5E, the gene encoding for CD73, is highly expressed in a MUC5B+ ductal cell population (**Figure 1A**). To confirm ductal cells expressed CD73, we used a human tissue microarray to evaluate cellularity of CD73 immunolabeling in chronic pancreatitis. An immunohistochemistry (IHC) stain for CD73 in 6 patient samples from human chronic pancreatitis (**Figure 1B**) revealed a strong positive staining in ADM cells (green arrows), ductal cells (red arrows) and infiltrating immune cells (yellow arrows).

**Figure 1.**
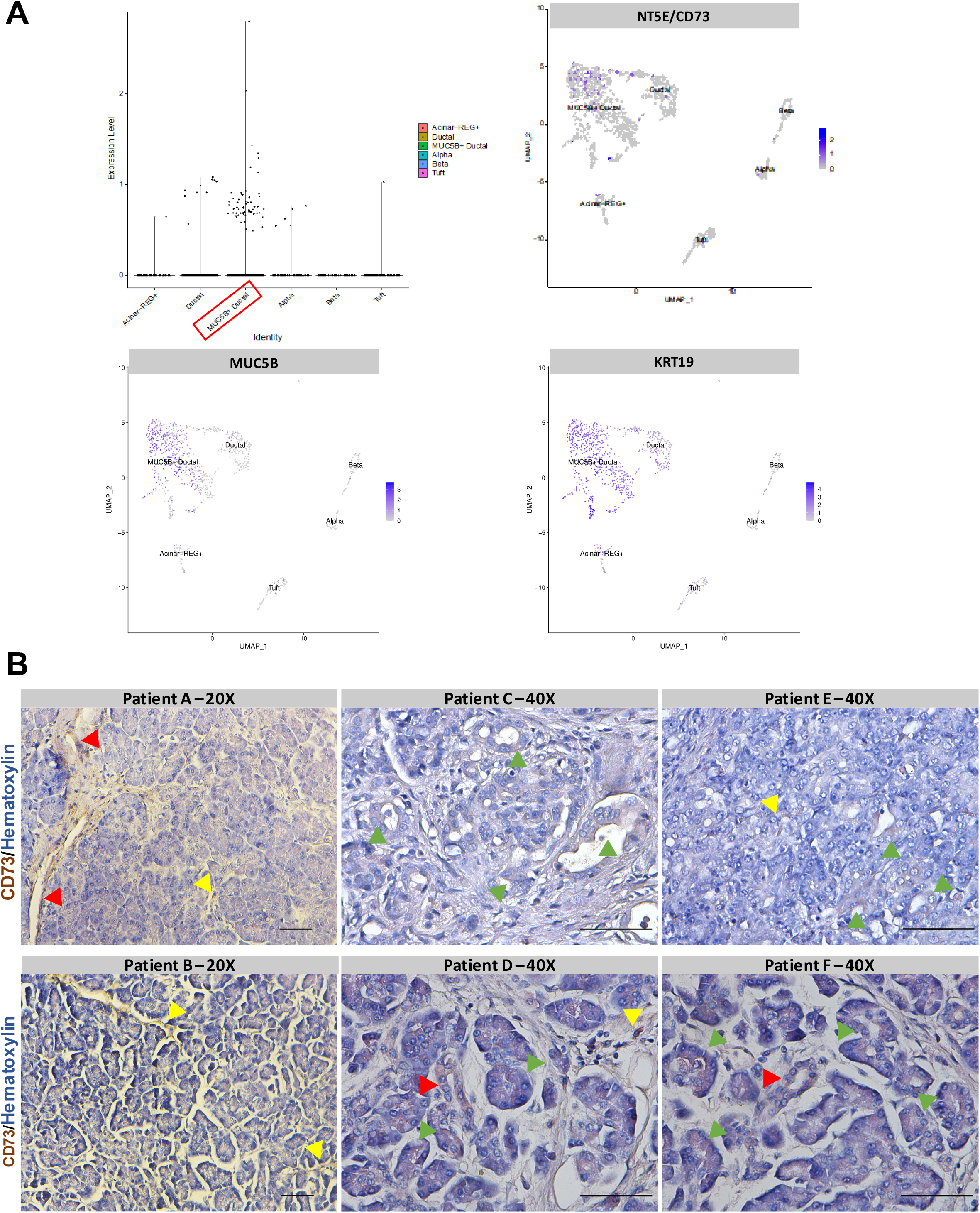
CD73 is expressed on ductal cells and infiltrating immune cells in human chronic pancreatitis. Single nuclear RNA sequencing of human chronic pancreatitis was analyzed. IHC for CD73 was performed in a TMA with 6 human cases of chronic pancreatitis. **A)** Single-nuclear RNA sequencing and associated UMAP from human chronic pancreatitis. NT5E (CD73) is expressed in a MUC5B+ ductal cell population (MUC5B+ and KRT19+ cells). **B)** Human chronic pancreatitis tissue (n=6) demonstrating positive CD73 staining in ADM cells (green arrows), ductal cells (red arrows) and infiltrating immune cells (yellow arrows). Bars represent 50uM.

### Genetic loss CD73 increases severity of chronic pancreatitis

To determine the role of CD73 in pancreatitis, we utilized a murine two-week chronic pancreatitis model (**Figure 2A**). Wild type and *CD73*^*-/-*^ mice were used for the study, and both genotypes were subjected to a caerulein-induced pancreatitis protocol consisting of two 250 ug/kg injections per day, 5 consecutive days a week, for two weeks. The mice were then euthanized after a two-day recovery period to determine expression of CD73 in murine pancreatitis and to evaluate if loss of CD73 resulted in any histopathologic changes in the pancreas compared to wild type pancreata. We determined the localization of CD73 *in vivo* by CD73 IHC staining and observed a strong positive expression of CD73 on infiltrating immune cells as well as ductal cells only in wild type mice under chronic pancreatitis, whereas the absence of CD73 expression was confirmed in pancreata from the in *CD73*^*-/-*^ mice (**Figure 2B, black arrows**).

**Figure 2.**
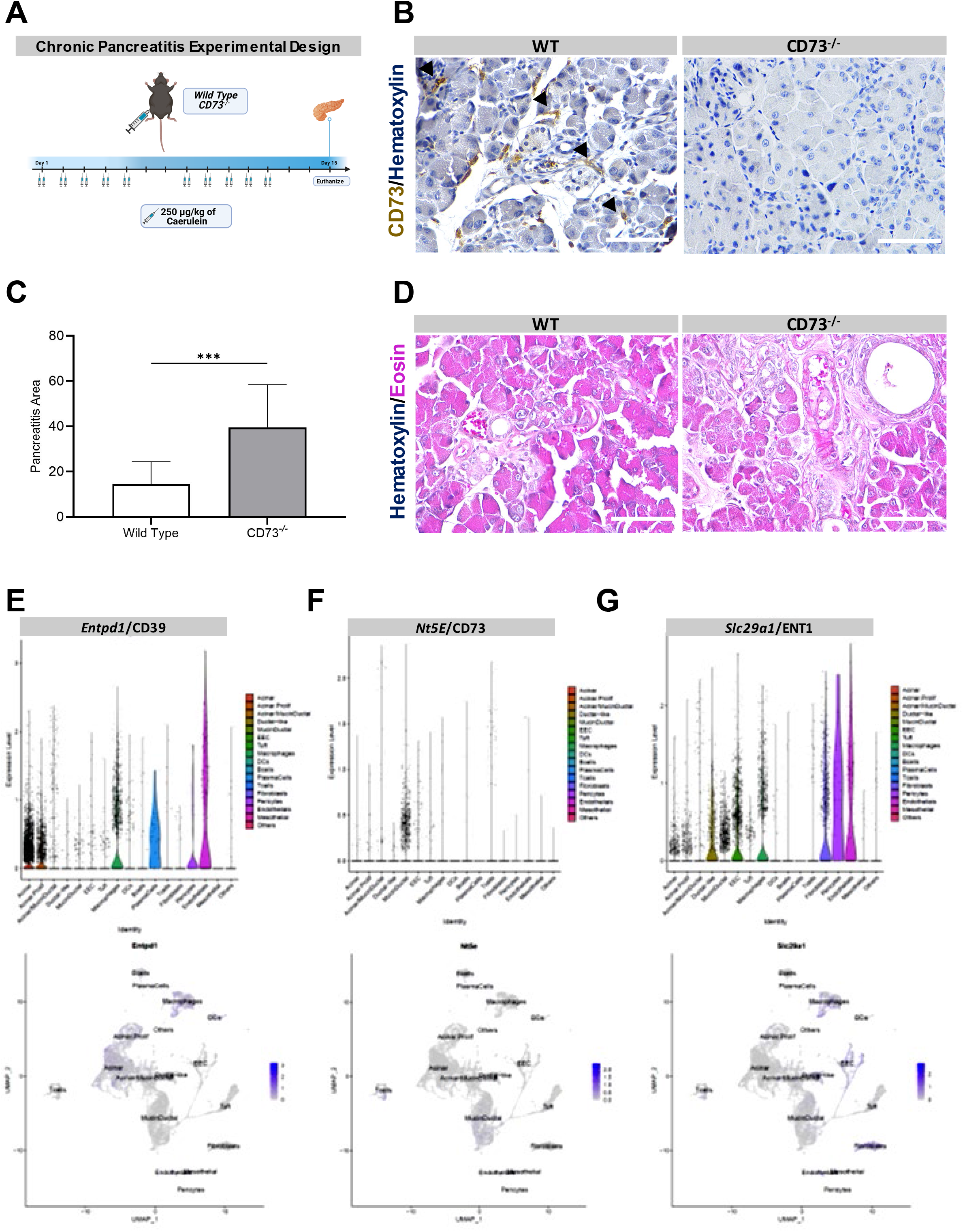
Genetic loss of CD73 increases severity of chronic pancreatitis. An *in vivo* model of chronic pancreatitis was performed in WT and *CD73*^*-/-*^ mice. Histopathology and purinergic enzymes expression were analyzed. **A)** Experimental design for the chronic pancreatitis model *in vivo*. **B)** IHC stain for CD73 in WT and *CD73*^*-/-*^ mice demonstrating positive staining on infiltrating immune cells as well as ductal cells only in WT mice (black arrows). **C)** ImageJ quantification of severe chronic pancreatitis revealed significantly increased severe pancreatitis area *CD73*^*-/-*^ mice compared to WT. Data was analyzed by Student’s t test. *D)* H&E stain of chronic pancreatitis in WT and *CD73*^*-/-*^ mice. **E-G)** Single-cell RNA sequencing and associated UMAP from a chronic pancreatitis mouse model at 2 and 4 weeks showed CD39 is highly expressed in acinar cells and macrophages **(E)**, CD73 is highly expressed in mucin/ductal cell populations **(F)** and ENT1 adenosine transporter is highly expressed in ductal-like, endothelial, EEC cell populations and macrophages **(G)**. Scale bars 50 μm. Error bars, SEM. ***□P□≤ 0.001.

To evaluate tissue injury and pathology in the context of chronic caerulein treatment, we used Hematoxylin & Eosin (H&E) to stain wild type and *CD73*^*-/-*^ mice. For comparison, severe pancreatitis was quantified and defined as the presence of significant infiltrating immune cells, ADM cells, and loss of normal cellular histology. Under caerulein-mediated chronic pancreatitis conditions, *CD73*^*-/-*^ mice displayed significantly increased severe pancreatitis area per field (*p < 0.001*) (**Figure 2C-D**) suggesting the loss of extracellular adenosine generation exacerbates and sustains tissue injury as well as inhibits tissue regeneration.

As we observed such a prominent difference in pancreatic injury in wild type compared to *CD73*^*-/-*^ mice after chronic injection of caerulein, we wanted to evaluate the cellular expression of CD73, CD39 and ENT1 in caerulein mediated murine chronic pancreatitis. We analyzed single-cell RNA sequencing data from a chronic caerulein-mediated mouse model recently published by Ma, et. al. encompassing ∼21,140 cells from 4 mice [51]. The results demonstrated the enzyme CD39, responsible for catalyzing the conversion of ATP to ADP and AMP, is highly expressed on macrophages, pericytes, endothelial cells and acinar cells (**Figure 2E**); whereas, similar to what we identified in human chronic pancreatitis, CD73 is highly expressed in a mucin/ductal cell population as well as in T cells, macrophages and B cells (**Figure 2F**). Lastly, the ENT1 adenosine transporter that facilitates the movement of extracellular adenosine across the cell membrane is highly expressed in macrophages, fibroblasts, pericytes, enteroendocrine cells, endothelial cells, and ductal-like cells (**Figure 2G**).

### Purinergic signaling modulates response to acute pancreatitis

As we observed such a significant difference in pancreatitis area in the chronic pancreatitis model, we wanted to evaluate the role of CD73 in a caerulein mediated acute pancreatitis model, which allows for histologic visualization of pancreatic repair over a time frame of 7 days after acute injury [55]. Wild type and *CD73*^*-/-*^ mice were used for the study and underwent a caerulein-induced acute pancreatitis protocol which consisted of eight injections of 70 ug/kg caerulein per day for two consecutive days (**Figure 3A**). Mice were then euthanized at 1, 4, and 7 days after the last caerulein injection to evaluate the timing of AMP and adenosine generation, which were also compared to chronic exposure to caerulein levels. Under acute pancreatitis, high performance liquid chromatography (HPLC) revealed AMP levels acutely decrease from Day 1 to Day 4, and increase from Day 4 to Day 7 in wild type mice; in contrast, in *CD73*^*-/-*^ mice these levels increase from Day 1 to Day 4 and decrease from Day 4 to Day 7, suggesting a transient accumulation of AMP in the context of no CD73 activity (**Figure 3B**). In wild type mice, acute pancreatitis showed AMP variations that were accompanied by a significant increase in ADO levels between Day 1 and Day 4, followed by a decrease between Day 4 and Day 7; however, no significant variations were observed in *CD73*^*-/-*^ mice (**Figure 3C**). Chronic exposure to caerulein showed increased AMP levels in *CD73*^*-/-*^ mice but decreased ADO levels when compared to wild type pancreata.

**Figure 3.**
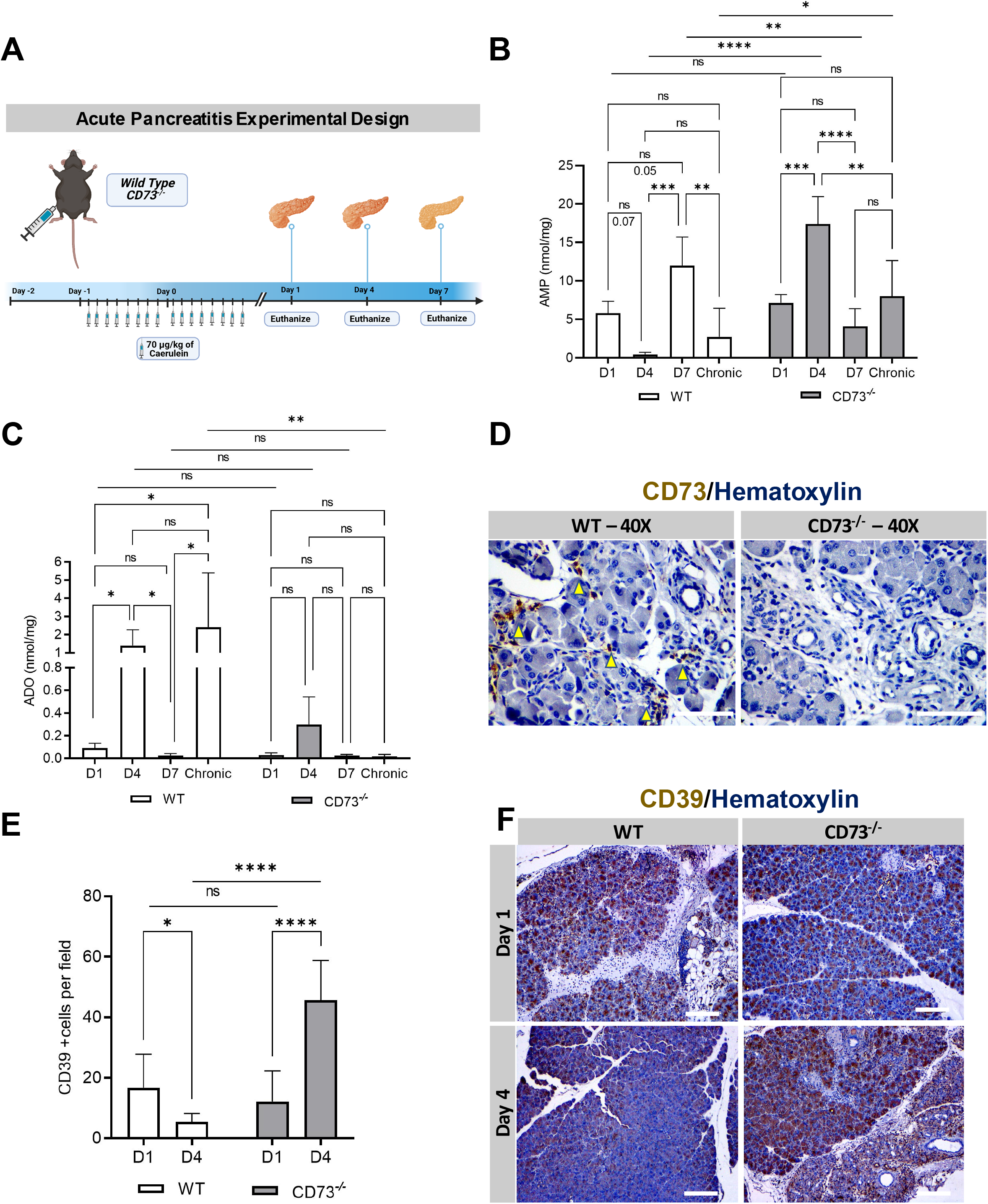
Purinergic signaling modulates response to acute pancreatitis. AMP and ADO levels were analyzed by HPLC from pancreatic tissues for AMP in WT mice at Days 1 (n = 4), 4 (n = 4), 7 (n = 3) and chronic model (n = 5) and *CD73*^*-/-*^ mice at Days 1 (n = 5), 4 (n = 4), 7 (n = 4), and chronic model (n = 5). CD39 and CD73 expression were analyzed by IHC. **A)** Experimental design for the acute□pancreatitis model *in vivo*. **B)** AMP levels acutely decrease from Day 1 to Day 4 and increase from Day 4 to Day 7 in WT mice; in contrast, in *CD73*^*-/-*^ mice these levels increase from Day 1 to Day 4 and decrease from Day 4 to Day 7. Chronic exposure to caerulein, showed increased AMP levels in *CD73*^*-/-*^ mice. **C)** ADO levels increased in WT mice between Day 1 and Day 4, followed by a decrease between Day 4 and Day 7; however, no significant variations were observed in *CD73*^*-/-*^ mice. Chronic exposure to caerulein, showed decreased ADO levels when compared to WT group. D1 and D4 were compared with Student’s t test. **D)** CD73 at Day 4 revealed positive staining on infiltrating immune cells (yellow arrows) in the WT only. **E)** ImageJ quantification of CD39 showed decreased expression between Day 1 and Day 4 in WT mice and increased in *CD73*^*-/-*^ murine pancreata. Additionally, at Day 4 a significant increase in CD39 expression was observed in *CD73*^*-/-*^ compared to the WT pancreata. **F)** IHC stain for CD39 in WT and *CD73*^*-/-*^ mice at Day 1 and 4. Data was analyzed by 2-way ANOVA. Error bars, SEM. *,□P□≤ 0.05; **□P□≤ 0.01; ***□P□≤ 0.001; **** P□≤ 0.0001;□n.s., not significant. Scale bars 50 μm.

ADO levels were significantly elevated at Day 4 during acute pancreatitis so we wanted to determine the localization of CD73 at Day 4. IHC showed CD73 expression in infiltrating immune cells as well as ductal cells (**Figure 3D, yellow arrows**) in wild type mice. CD73 staining was also performed on *CD73*^*-/-*^ mice to confirm the genotype, which correctly demonstrated negative staining of the tissue.

To compare the initial modulation in nucleotide generation under acute conditions in wild type and *CD73*^*-/-*^ mice, an IHC antibody stain for CD39 was performed at Day 1 and Day 4. CD39 expression decreased between Day 1 and Day 4 in wild type mice and increased in *CD73*^*-/-*^ murine pancreata (**Figure 3E-F**). Additionally, at Day 4 a significant increase in CD39 expression was observed in *CD73*^*-/-*^ compared to the wild type pancreata. These data support the HPLC data that *CD73*^*-/-*^ mice are experiencing enhanced nucleotide or purinergic signaling at Day 4, which may be a major determinant of sustained tissue injury in *CD73*^*-/-*^ mice, while the wild type mice utilize CD73 to convert AMP to adenosine.

### Genetic loss of CD73 promotes metaplasia in acute pancreatitis

In order to better comprehend the initial tissue injury response and resolution in the acute pancreatitis model, we evaluated immune infiltration and metaplasia. To correlate tissue injury to the amount of ADM cells present, an IHC antibody stain for Cytokeratin-19, a marker for ductal cells, was performed in wild type and *CD73*^*-/-*^ mice at Day 1 and Day 4 (**Figure 4A**). At Day 1, the number of ductal cells per field between experimental groups was similar, indicating comparable initial tissue injury and metaplasia (*p = n.s*.) (**Figure 4B**). However, at Day 4, *CD73*^*-/-*^ mice displayed a significant increase in the amount of Cytokeratin-19+ areas per field, indicating a significant increase in metaplastic ducts and ADM, compared to wild type pancreata (*p < 0.001*). Interestingly, from Day 1 to Day 4 there was a significant increase in Cytokeratin-19+ cells in both wild type mice (*p < 0.01*) and *CD73*^*-/-*^ mice (*p < 0.0001*) (**Figure 4B**). These findings indicate the ADM process is a reparative mechanism concurrent with peak pancreatic adenosine generation.

**Figure 4.**
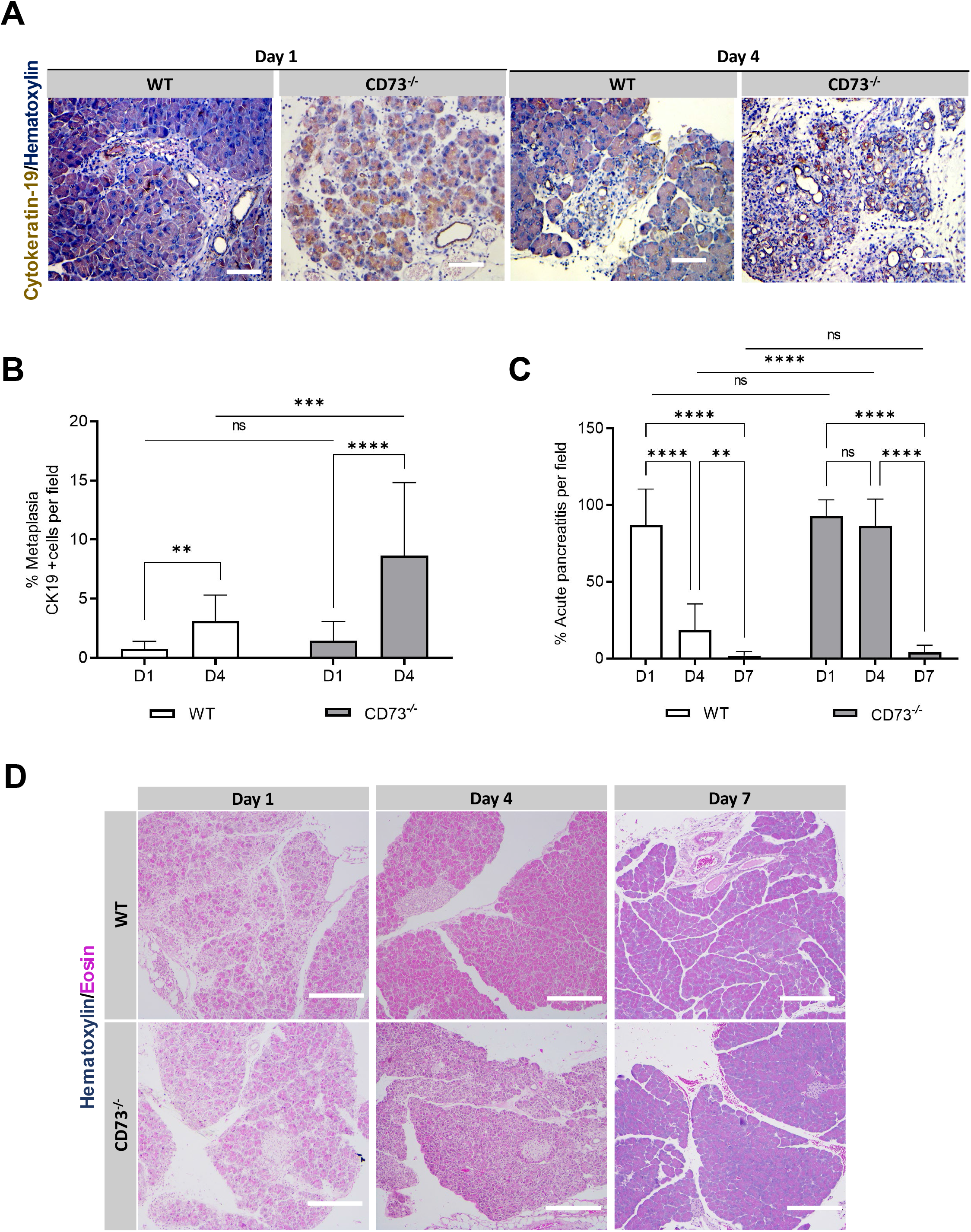
Genetic loss of CD73 promotes metaplasia in acute pancreatitis. Metaplasia was studied by Cytokeratin 19 IHC staining in WT and *CD73*^*-/-*^ mice at days 1 and 4. Pancreatitis area was evaluated in WT mice at Days 1 (n = 12) and 4 (n = 14) and in *CD73*^*-/-*^ mice at Days 1 (n = 12) and 4 (n = 21). **A)** IHC stain for Cytokeratin-19 in WT and *CD73*^*-/-*^ mice at Days 1 and 4. **B)** ImageJ quantification of Cytokeratin-19 positive areas per field a significant increase from Day 1 to Day 4 in both genotypes. Additionally, at Day 4, the presence of Cytokeratin-19+ cells were significantly increased in *CD73*^*-/-*^ mice compared to WT. Student’s t test was used to compare WT timepoints. **C)** ImageJ quantification of acute pancreatitis revealed increased persistent pancreatitis in *CD73*^*-/-*^ mice compared to WT. By Day 7, both genotypes demonstrated similar histology with no significant difference in pancreatitis. □**D)**□H&E stain of acute pancreatitis in wild type and *CD73*^*-/-*^ mice at Days 1, 4, and 7. Data was analyzed by 2-way ANOVA. Error bars, SEM. **□P□≤ 0.01; ***□P□≤ 0.001; **** P□≤ 0.0001;□n.s., not significant. Scale bars 50 μm.

To evaluate severity of pancreatitis at Day 4 and Day 7, an H&E stain was performed in wild type and *CD73*^*-/-*^ mice at Day 1, 4, and 7. At Day 1, both experimental groups showed increased fluid between the pancreatic lobes and acinar cells, similar presence of ADM cells, and infiltrating immune cells (**Figure 4C-D**). At Day 4, wild type mice demonstrated a return to normal histology characterized by the disappearance of ADM cells and infiltrating immune cells and reduction in excess fluid, while *CD73*^*-/-*^ mice showed significantly increased residual pancreatitis areas (*p < 0.0001*). Lastly, at Day 7 both experimental groups demonstrated near complete return to normal histology with no significant difference between them (*p = n.s*.). These data suggest *CD73*^*-/-*^ mice exhibit sustained tissue injury and require more time for tissue regeneration compared to the wild type mice.

### Loss of adenosine increases immune infiltration in acute pancreatitis

To evaluate mediators of sustained inflammation in the *CD73*^*-/-*^ mice, we used IHC to stain for Granzyme B, Myeloperoxidase (MPO) and NIMPR14, a neutrophil marker. IHC experiments were performed in wild type and *CD73*^*-/-*^ mice at Day 1 and Day 4. At Day 1, the number of Granzyme B+ cells were similar between experimental groups (*p = n.s*.) (**Figures 5A and 5D**). However, at Day 4 *CD73*^*-/-*^ mice demonstrated significantly increased Granzyme B+ cells compared to the wild type mice (*p < 0.0001*). Additionally, from Day 1 to Day 4 there was a significant increase in Granzyme B+ cells in *CD73*^*-/-*^ mice (*p < 0.05*) and a significant decrease in the wild type (*p < 0.05*). These data suggest Granzyme B expressing cells are a major determinant of pancreatitis severity in the absence of extracellular adenosine.

**Figure 5.**
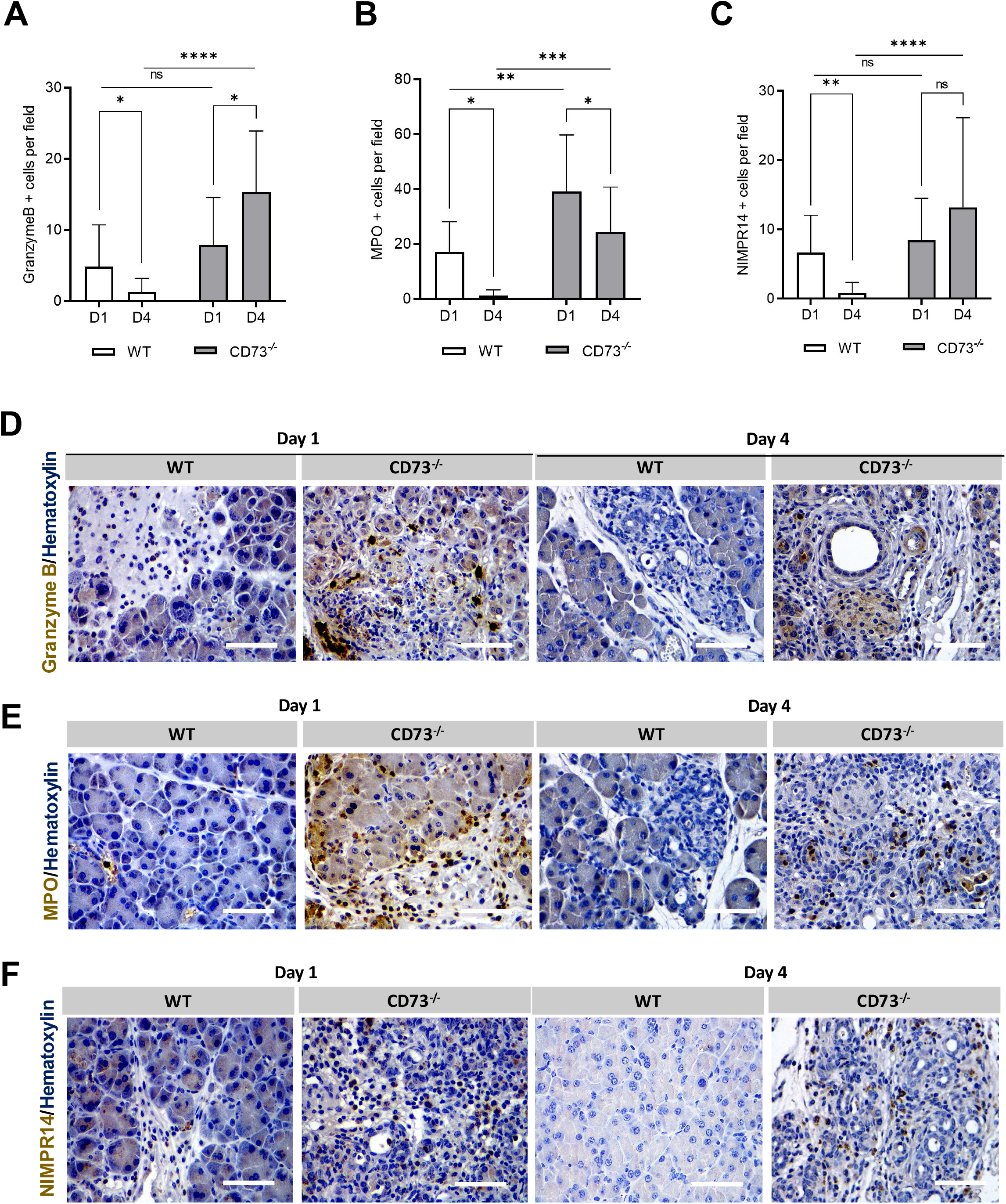
Loss of adenosine increases immune infiltration in acute pancreatitis. Immune cell infiltration was analyzed by IHC in WT and *CD73*^*-/-*^ mice at days 1 and 4. **A)** ImageJ quantification of Granzyme B+ cells per field, a marker for T-cells and Natural Killer cells, was increased only in *CD73*^*-/-*^ mice at D4 when compares with D1 and WT D4 measurements. WT mice presented decreased Granzyme B + cells at D4 compared to D1. Student’s t test was used to analyze WT timepoints. **B)** ImageJ quantification of MPO+ cells per field showed a decrease from D1 to D4 in both genotypes. Additionally, *CD73*^*-/-*^ mice presented increased MPO positive cells at both timepoint when compared to WT mice. **C)** ImageJ quantification of NIMPR-14+ cells showed significantly increased staining in *CD73*^*-/-*^ mice when compared to WT. In WT mice, levels decreased from D1 to D4. Student’s t test was used to analyze WT timepoints. Representative images for Granzyme B **(D)**, NIMPR14 **(E)** and NIMPR14 **(F)** IHC staining. Data was analyzed by 2-way ANOVA. Error bars, SEM. *,□P□≤ 0.05; **□P□≤ 0.01; ***□P□≤ 0.001; **** P□≤ 0.0001;□n.s., not significant. Scale bars 50 μm.

To determine the innate immune system’s role in initial tissue injury and resolution, an IHC stain for MPO, a marker for inflammatory neutrophils, was performed in wild type and *CD73*^*-/-*^ mice at Day 1 and Day 4 (**Figure 5B and 5E**). From Day 1 to Day 4 there was a significant decrease in MPO+ cells in both wild type and *CD73*^*-/-*^ mice (*p < 0.05*). Interestingly, *CD73*^*-/-*^ mice demonstrated a significantly increased amount of MPO+ cells per field compared to the wild type mice at both Day 1 and Day 4 measurements (*p < 0.01*; *p < 0.001*, respectively), suggesting the absence of extracellular adenosine in *CD73*^*-/-*^ mice promotes early tissue injury via innate immune cell infiltration.

Due to the potent induction of the innate immune system at Day 1 as shown in the MPO IHC, we decided to specifically evaluate neutrophil activity. An IHC stain for NIMPR-14, a marker for Ly6G+ and Ly6C+ neutrophils, was performed in wild type and *CD73*^*-/-*^ mice at Day 1 and Day 4 (**Figure 5C and 5F**). At Day 1, the number of neutrophils per field was similar between experimental groups (*p = n.s*.); however, at Day 4 *CD73*^*-/-*^ mice demonstrated significantly increased neutrophils per field compared to the wild type (*p < 0.0001*). From Day 1 to Day 4 there was a significant decrease in neutrophils in wild type mice (*p < 0.01*); while no difference was observed in neutrophil abundance in the *CD73*^*-/-*^ (*p = n.s*.). These results suggests that the sustained tissue injury seen in *CD73*^*-/-*^ mice at Day 4 is primarily due to the continued induction and activation of neutrophils in the absence of extracellular adenosine and possibly due to heightened ATP or AMP-dependent purinergic signaling.

### Adenosine restrains neutrophil-mediated tissue injury

To confirm that adenosine generation promoted neutrophil-mediated tissue injury, we modified the caerulein induced acute pancreatitis model by treating *CD73*^*-/-*^ mice with a Ly6G neutrophil depletion antibody or vehicle (**Figure 6A**) and euthanized the mice at Day 4 post last caerulein injection. To confirm neutrophil depletion, an IHC stain for NIMPR-14 was performed which demonstrated the presence of neutrophils in the vehicle treated and the absence of neutrophils in pancreata from the neutrophil depleted mice (**Figure 6B**). We performed a H&E stain to assess tissue injury, and found that at Day 4, neutrophil depleted *CD73*^*-/-*^ mice demonstrated significantly less pancreatitis per field compared to vehicle treated (*p < 0.05*) (Figures 6C-D), indicating enhanced neutrophil activity from loss of CD73 dependent adenosine generation promotes sustained inflammation.

**Figure 6.**
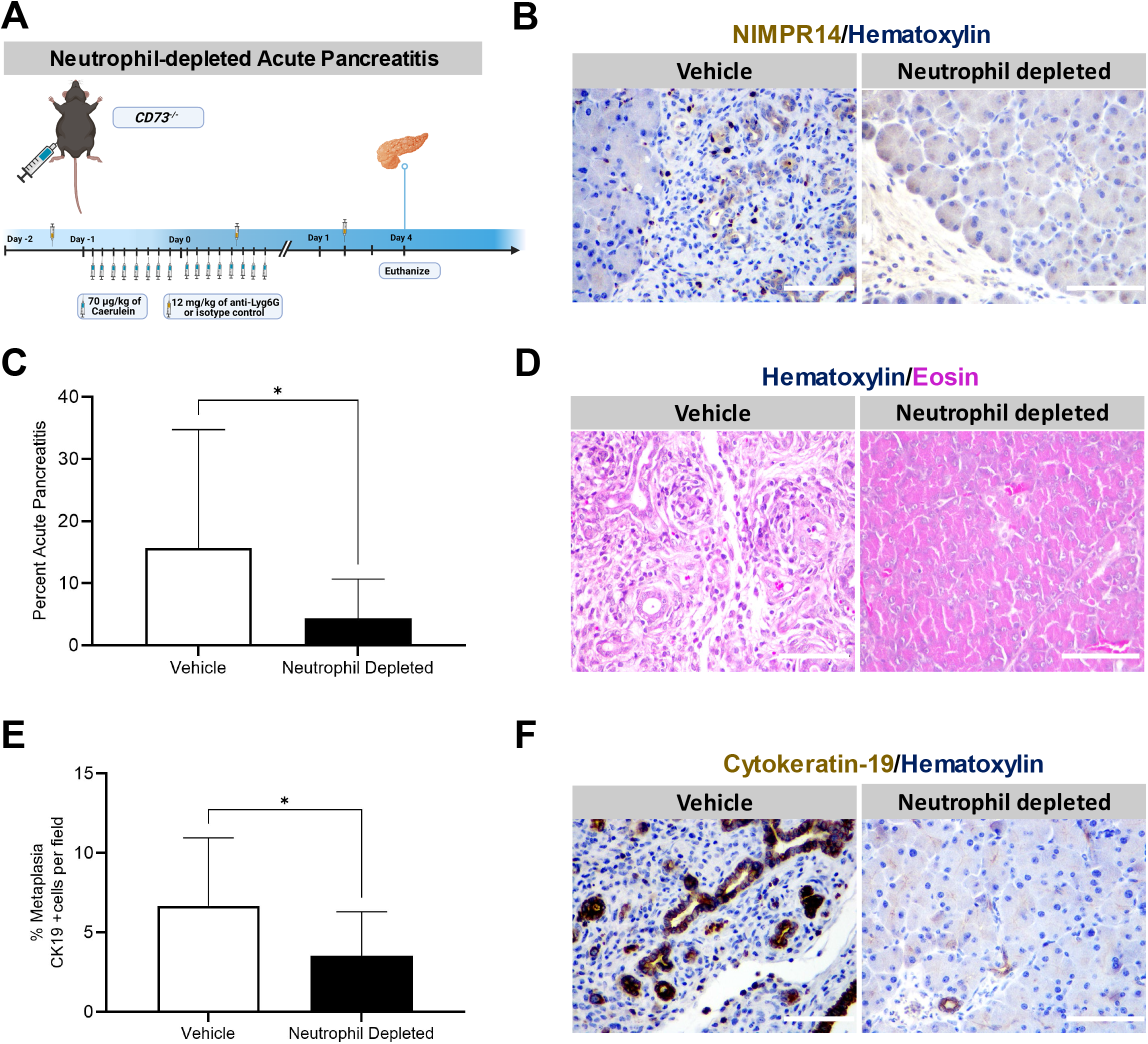
Genetic loss of adenosine increases neutrophil-mediated oxidative stress-induced tissue injury. A neutrophil depletion *in vivo* experiment was performed in WT and *CD73*^*-/-*^ mice subjected to acute pancreatitis. Mice were euthanized at Day 4. Pancreatitis and cytokeratin 19 areas were assessed by histopathology and IHC. **A)** Experimental design for the neutrophil-depleted acute pancreatitis model *in vivo*. **B)** IHC stain for NIMPR-14 was performed in vehicle treated and neutrophil depleted *CD73*^*-/-*^ mice at Day 4 to confirm depletion. **C)** ImageJ quantification at day 4 revealed decreased pancreatitis area per field in neutrophil depleted mice compared to vehicle treated. **D)** H&E stain in neutrophil depleted acute pancreatitis mice model at Day 4. **E)** ImageJ quantification of Cytokeratin-19 positive areas per showed decreased levels in neutrophil depleted mice compared to vehicle treated. **F)** IHC stain for Cytokeratin-19 in neutrophil depleted acute pancreatitis mice model at Day 4. Data was analyzed by Student’s t test. Error bars, SEM. *,□P□≤ 0.05. Scale bars 50 μm.

To determine if neutrophil-depletion could prevent metaplasia in acute pancreatitis, an IHC stain for Cytokeratin-19 was performed in vehicle treated and neutrophil depleted *CD73*^*-/-*^ mice at Day 4, which showed vehicle treated mice, compared to neutrophil depleted mice, presents significantly increased amounts of metaplasia per field, suggesting the absence of neutrophils restrains metaplasia in acute pancreatitis (*p < 0.05*) (**Figures 6E-F**).

### Enhanced adenosine receptor activation promotes resolution of tissue injury in acute pancreatitis

To determine if enhanced adenosine receptor activation would reduce tissue injury and metaplasia, wild type and *CD73*^*-/-*^ mice were administered NECA, a high affinity adenosine receptor agonist, in an acute pancreatitis model (**Figure 7A**). We assessed tissue injury in H&E staining and found *CD73*^*-/-*^ mice demonstrate significantly less pancreatitis area per field compared to their caerulein-only treated genotype comparison (*p < 0.01*) (**Figures 7B-C**). To evaluate the effect of enhanced adenosine generation on metaplasia pancreatitis area, an IHC stain for Cytokeratin-19 was conducted to investigate the amount of metaplastic ductal cells present at Day 1 in NECA-treated wild type and *CD73*^*-/-*^ mice. Our analysis revealed NECA-treated *CD73*^*-/-*^ mice present significantly decreased amount of metaplasia compared to the caerulein-only treated *CD73*^*-/-*^ mice (*p < 0.05*) (**Figures 7D-E**). In contrast, there were similar levels of Cytokeratin-19 positive cells in NECA-treated and caerulein-only treated wild type mice at Day 1 (*p = n.s*.), suggesting that the most significant therapeutic effect was seen in animals that were previously adenosine depleted.

**Figure 7.**
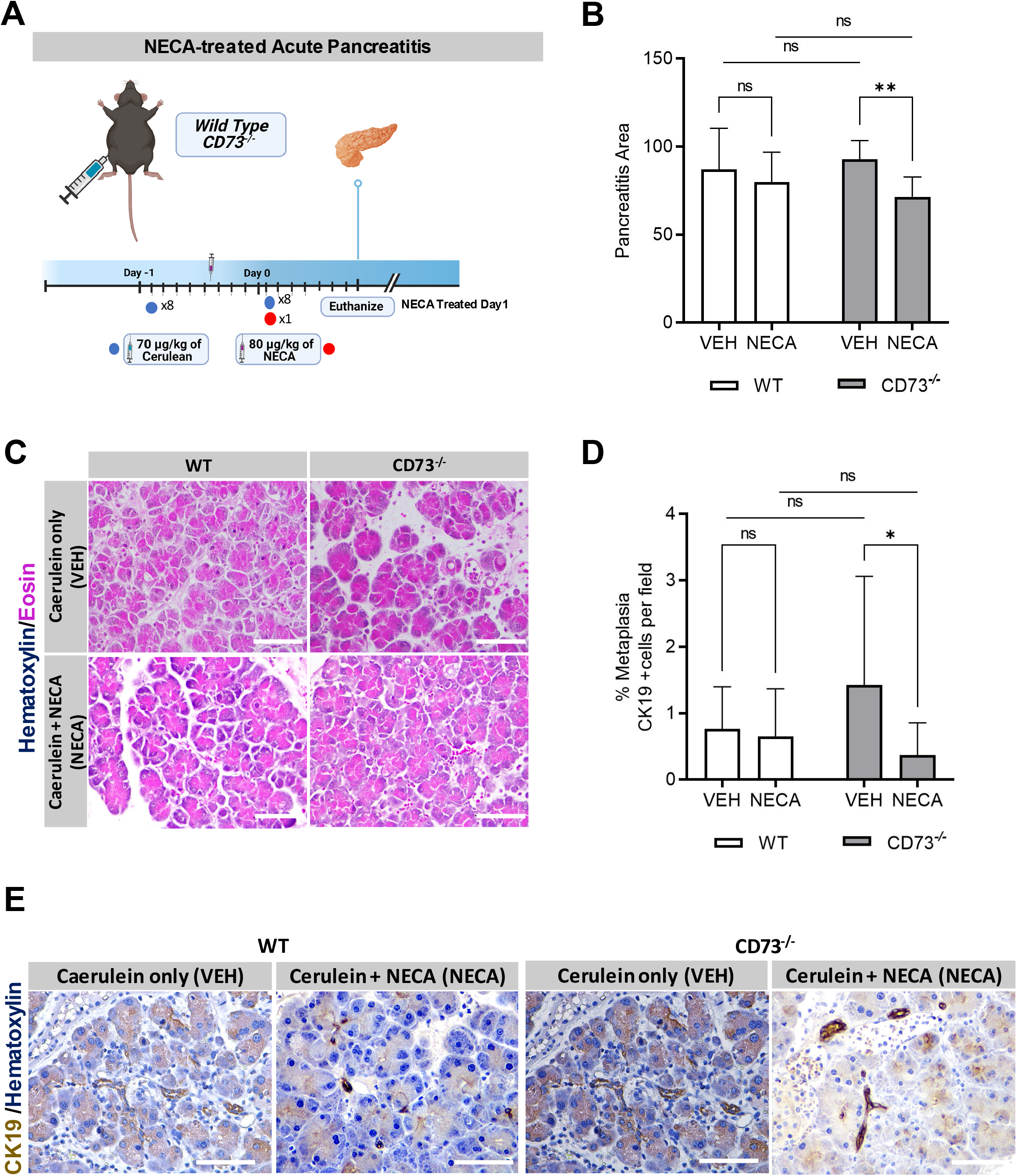
Enhanced adenosine receptor activation promotes resolution of tissue injury and metaplasia in acute pancreatitis. An acute pancreatitis *in vivo* experiment was performed in *CD73*^*-/-*^ mice treated with NECA or vehicle. Pancreatitis and Cytokeratin 19+ areas were assessed by histopathology and IHC. **A)** Experimental design for the NECA-treated acute pancreatitis model *in vivo*. **B)** ImageJ quantification of pancreatitis per field revealed NECA treated *CD73*^*-/-*^ presented less pancreatitis area than vehicle treated mice after acute pancreatitis induction. **C)** H&E stain of VEH and NECA treated WT and *CD73*^*-/-*^ mice at Day 1 in the acute pancreatitis model. **D)** ImageJ quantification of Cytokeratin-19 positive areas per field showed increased metaplastic ductal cells per field in caerulein-only treated *CD73*^*-/-*^ mice compared to NECA treated. **E)** IHC stain for Cytokeratin-19 in VEH and NECA treated WT and *CD73*^*-/-*^ mice at Day 1. Data was analyzed by 2-way ANOVA. Error bars, SEM. *,□P□≤ 0.05; **□P□≤ 0.01;□n.s., not significant. Scale bars 50 μm.

## DISCUSSION

Acinar and ductal cells comprise the exocrine component of pancreatic parenchymal function. Acinar cells are highly specialized cells characterized by zymogen granules and abundant rough endoplasmic reticulum. Acinar cells are responsible for synthesizing, storing and secreting digestive enzymes including amylase, lipase, peptidase and nucleases. In addition, acinar cells are the major source of trypsinogen, a component of pancreatic juice that is cleaved to trypsin by enteropeptidases in the intestinal mucosa [56, 57]. The main function of the pancreatic ducts is to carry fluid containing digestive enzymes secreted from acinar cells. Ductal cells are a major source of sodium bicarbonate (NaHCO3) rich fluid which flushes out and neutralizes pH from digestive enzymes [58]. In pancreatitis, the ductal epithelium does not appropriately regulate pH and more neutral pH or acidic pH is generated, causing obstruction and dilation of the duct lumen [59]. The ductal system is intricate, with many small peripheral ducts all channeling to the main pancreatic duct. When gallstones, calcification or intraductal lesions block the exocrine component of the pancreas, patients develop pain and inflammation as there is abnormal release of digestive enzymes, which can contribute to pancreatic fibrosis, calcification and downstream pathophysiology. Neutrophil infiltration is one of the first pathogenic responses in early phases of pancreatitis [60] and neutrophil accumulation is thought to prematurely activate trypsinogen release and aid in progression from acute to severe acute and chronic pancreatitis through production of reactive oxygen species (ROS) and hydrolases (reviewed in [60]). Neutrophil depletion significantly reduces serum amylase and reduces pancreatic injury in models of severe acute pancreatitis [61-63]. The coordinated role of innate immune cells, including macrophages, and intrapancreatic cytokines and chemokines in pancreatic injury is important for resolution of injury and restoration of organ function.

In this manuscript, we describe that ductal cells and possibly subsets of ADM can express CD73 in the context of caerulein-mediated acute or chronic pancreatitis. Through analysis of published single nuclear or single cell RNA-seq datasets and immunohistochemistry, we show ductal cells and immune cells within interstitial spaces that express CD73 and we demonstrate acinar-to-ductal metaplasia may be a CD73-mediated reparative process in the pancreas. We show CD73 promotes adenosine generation, a critical nucleoside to resolve tissue injury in response to pancreatitis and suggest the increased acinar-to-ductal metaplastic cells seen in *CD73*^*-/-*^ mice during acute pancreatitis may arise as an attempted tissue repair process. Under chronic conditions we observe expression of CD39 *in vivo* on macrophages which may indicate that the innate immune system is contributing to sustained injury, while the relatively high expression of CD73 *in vivo* on immune cells including T cells may indicate a divergent immune-mediated mechanism is contributing to CD73-mediated tissue injury resolution. Future studies delineating the role of specific adenosine receptors in stromal and immune cells in pancreatitis models will establish the mechanistic consequences of elevated intrapancreatic adenosine on development and resolution of pancreatitis.

Sustained enzymatic activity of CD39 in *CD73*^*-/-*^ mice implicates enhanced AMP and loss of extracellular adenosine which contribute to enhanced disease severity in pancreatitis. Extracellular AMP is necessary for the initial response to acute injury in the pancreas by promoting inflammation, as seen by similar levels of CD39 in both wild type and *CD73*^*-/-*^ mice at Day 1. However, sustained AMP levels promotes a more severe phenotype. Thus, after initial insult, in wild type mice at Day 4, CD73 enzymatic activity is increased to promote adenosine generation and signaling through adenosine receptors, which promotes inflammation resolution. However, in *CD73*^*-/-*^ mice, sustained CD39 activity at Day 4 demonstrates reduced capacity to generate extracellular adenosine in the absence of CD73. This switch in modulation is also seen by HPLC in wild type mice by the upregulation of purinergic signaling at initial tissue injury and then significant downregulation of AMP levels at Day 4 while simultaneously promoting adenosine generation after sustained tissue injury to promote resolution.

While both genotypes showed similar tissue injury and histologic change at Day 1 in the acute pancreatitis model, by Day 4 the wild type mice showed a near complete resolution of inflammation, while the *CD73*^*-/-*^ mice demonstrated significant residual pancreatitis injury. The increased presence of acinar-to-ductal metaplastic cells at Day 4 in *CD73*^*-/-*^ mice compared to the wild type suggests that without adenosine generation, reparative processes are still necessary to mitigate pancreatic injury. Additionally, increased staining of Cytokeratin-19 in pancreatitis areas compared to normal histologic areas in both wild type and *CD73*^*-/-*^ mice, but even greater still in *CD73*^*-/-*^ mice, demonstrates the response to injury is regulated by adenosine receptor activation. This suggests the enhanced CD73 activity partially by ductal cells or ADM and most prominently by extra-epithelial cells in wild type mice is allowing for an immune-mediated rapid resolution of pancreatitis. However, at Day 7 in *CD73*^*-/-*^ mice there is also a near complete resolution of tissue injury. In addition, we observe by HPLC adenosine in *CD73*^*-/-*^ mice at Day 4 indicating intracellular conversion of AMP to adenosine and subsequent transport of adenosine into the microenvironment by ENT1 transporters may have been mechanistically why *CD73*^*-/-*^ mice were able to resolve caerulein-induced pancreatitis by Day 7. Further studies will be required to investigate cellularity of adenosine receptor expression to determine how adenosine specifically reduces pancreatic inflammation.

In these experiments, in addition to histological changes and differences in ADM abundance, we quantified a significant increase in MPO+ Ly6G+ and Granzyme B+ cells in *CD73*^*-/-*^ mice at Day 4, which demonstrates a potent pro-inflammatory response in the absence of adenosine in response to caerulein-induced injury. The binding of ATP to the P2X and P2Y families of receptors, expressed on neutrophils in humans and *in vivo* [45], enhances neutrophil phagocytosis, chemotaxis, and oxidative burst. Under conditions of enhanced purinergic signaling, the resulting sustained activity and chemotactic ability of neutrophils promote sustained tissue injury and slower inflammation resolution. We experimentally show the significant impact of neutrophils in our genetic model using neutrophil deletion experiments which significantly reduced inflammation in *CD73*^*-/-*^ mice. In addition, in NECA treated mice, we predict the high adenosine concentration at Day 1 promotes adenosine dependent activation of A_2A_ and A_2B_ adenosine receptors over the high affinity A_1_ receptor to promote tissue regeneration and restrain metaplasia in acute pancreatitis [35, 42]. This identifies CD73 agonists as a potential therapeutic target for patients with acute pancreatitis as a mechanism to rapidly eliminate persistent inflammation via activation of A_2A_ and A_2B_ receptors.

## METHODS

### Animal Model

All mouse model procedures are in compliance with UTHealth’s CLAMC Animal Welfare Committee Review and approved on Dr. Bailey-Lundberg’s AWC protocol. To evaluate the role of CD73 in pancreatitis, CD73 knockout (*CD73*^*-/-*^) and C57BL/6 (wild type) mice were used. Full body CD73KO mice were purchased from The Jackson Laboratory strain 018986. Controls for each experiment were derived from wild type crosses. Caerulein injections were performed for each experiment during Spring months (March-June) 2022 and mixed genders were equally included for all groups. In the acute pancreatitis model, *CD73*^*-/-*^ and wild type mice were intraperitoneally injected on alternating flanks with 70 μg/kg of caerulein (Sigma Aldrich 17650-98-5) 8x a day for two consecutive days. Mice were euthanized at 1, 4, and 7 days after the last caerulein injection to evaluate ADM abundance, inflammatory progression and organ repair. For the chronic model, *CD73*^*-/-*^ and wild type mice were injected intraperitoneally on alternating flanks with 250 μg/kg of caerulein (Sigma Aldrich 17650-98-5) twice a day, 5 days a week, for two weeks. The mice were allowed to recover for two days after the last injection before euthanasia by isoflurane overdose.

### Neutrophil Depletion *in vivo*

*CD73*^*-/-*^ mice were intraperitoneally administered 300 μg of anti-Ly6G antibody (clone1A8, BioXCell, West Lebanon, NH) at Day -2, 0, and 1 day after the last caerulein injection. IgG2a isotype (clone 2A3, BioXCell, West Lebanon, NH) was used for control. Mice were euthanized at Day 4.

### NECA *in vivo*

*CD73*^*-/-*^ and wild type mice under acute pancreatitis protocol of 8 injections per day during 2 consecutive days (Days -1 and 0) were administered a single injection of 80 μg/kg of 5’-N-Ethylcarboxamidoadenosine (NECA) (MedChemExpress HY-103173) after the sixth injection of caerulein on day zero. Mice were euthanized at Day 1. NECA injection was performed intraperitoneally.

### Immunohistochemistry and ImageJ analysis

Tissues were fixed in zinc buffered formalin, processed according to standard protocols, and embedded in paraffin. The unstained sections were baked at 60 °C for 45 minutes. The sections were deparaffinized with Histoclear and rehydrated stepwise. Heat-mediated antigen retrieval followed with a pH 6 unmasking solution (Vector Laboratories, H-3300) and a pH 9 unmasking solution (Abcam 100X Tris-EDTA). All sections were blocked for one hour in 10% FBS in PBST. Primary antibodies were used at a 1:200 dilution and incubated overnight at 4 °C. Secondary antibodies were used at a 1:500 dilution and incubated at room temperature for 30 minutes. The Vectastain ABC kit Peroxidase Standard (Vector Laboratories, PK4000) and DAB Peroxidase (HRP) Substrate kit (Vector Laboratories, SK-4100) were used.

Image analysis was performed using ImageJ (http://imagej.nih.gov/ij/) software and 3-5 representative fields per tissue were used depending on the size of the tissue. The color threshold tool was used to determine positive staining and ensure normalization of all samples. Freehold selection tool was used to isolate mild and severe pancreatitis from normal tissue. Pancreatitis areas were defined by the presence of swelling between pancreatic lobes, significant infiltrating immune cells, ADM cells, and loss of normal cellular histology. Human pancreatitis tissues were obtained from a TMA array (US Biomax, Inc. BIC14011b).

### High Performance Liquid Chromatography (HPLC)

Pancreas tissue was collected at the end of the experiment and flash frozen. Samples were analyzed by high-performance liquid chromatography using the Waters Breeze 2 HPLC System (Waters 2489 UV/Visible Detector and Waters 1525 Binary HPLC Pump). Flow rate was 1 mL/minute and 100 uL per sample was injected. Absorbance was measured at a wavelength of 260 nm and 280 nm, and adenosine and AMP peaks were determined using a standard HPLC curve. Pancreas tissue adenosine and AMP levels were normalized to lysate protein levels.

#### Analysis of published single cell RNA sequencing datasets

Processed count matrices for scRNA-seq datasets from Ma et al. were downloaded from the Gene Expression Omnibus (GEO) database (accession number GSE172380). The processed human pancreas sNuc-seq dataset from Tosti et al. was obtained from http://singlecell.charite.de/pancreas/[64]. Low-quality cells were filtered on read counts, the number of genes expressed, and the ratio of mitochondrial reads following the thresholds described in the respective publications. Filtered gene count matrices were log-normalized, and the top 2000 variable features were further scaled prior to dimension reduction by PCA and being embedded in UMAP using the R package Seurat[65]. Seurat cell clusters were labeled with major cell types using marker genes provided by the authors.

## Abbreviations

ADM: acinar-to-ductal metaplasia
AMP: adenosine monophosphate
ATP: adenosine triphosphate
CCK: cholecystokinin
CD39: ectonucleotidase triphosphate diphosphohydrolase-1
EEC: enteroendocrine
HPLC: high performance liquid chromatography
IHC: immunohistochemistry
TNF: tumor necrosis factor α
NECA: 5’-N-Ethylcarboxamidoadenosine

